# Using photographs and deep neural networks to understand flowering phenology and diversity in mountain meadows

**DOI:** 10.1101/2023.03.28.533305

**Authors:** Aji John, Elli J. Theobald, Nicoleta Cristea, Amanda Tan, Janneke Hille Ris Lambers

**Affiliations:** Department of Biology, University of Washington, Seattle, WA 98195, USA; Department of Civil and Environmental Engineering, University of Washington, Seattle, WA 98195, USA; eScience Institute, University of Washington, Seattle, WA 98195, USA; Plant Ecology, Institute of Integrative Biology, D-USYS, ETH Zürich, 8092 Zürich, Switzerland

**Keywords:** Alpine wildflowers, convolutional neural net, climate-change, phenology

## Abstract

Mountain meadows are an essential part of the alpine-subalpine ecosystem; they provide ecosystem services like pollination and are home to diverse plant communities. Changes in climate affect meadow ecology on multiple levels, for example by altering growing season dynamics. Tracking the effects of climate change on meadow diversity through the impacts on individual species and overall growing season dynamics is critical to conservation efforts. Here, we explore how to combine crowd sourced camera images with machine learning to quantify flowering species richness across a range of elevations in alpine meadows located in Mt Rainier National Park, Washington, USA. We employed three machine learning techniques (Mask R-CNN, RetinaNet and YOLOv5) to detect wildflower species in images taken during two flowering seasons. We demonstrate that deep learning techniques can detect multiple species, providing information on flowering richness in photographed meadows. The results indicate higher richness just above the tree line for most of the species, which is comparable with patterns found using field studies. We found that the two-stage detector Mask R-CNN was more accurate than single-stage detectors like RetinaNet and YOLO, with the Mask R-CNN network performing best overall with mean average precision (mAP) of 0.67 followed by RetinaNet (0.5) and YOLO (0.4). We found that across the methods using anchor box variations in multiples of 16 led to enhanced accuracy. We also show that detection is possible even when pictures are interspersed with complex backgrounds and are not in focus. We found differential detection rates depending on species abundance, with additional challenges related to similarity in flower characteristics, labeling errors, and occlusion issues. Despite these potential biases and limitations in capturing flowering abundance and location-specific quantification, accuracy was notable considering the complexity of flower types and picture angles in this data set. We therefore expect that this approach can be used to address many ecological questions that benefit from automated flower detection, including studies of flowering phenology and floral resources, and that this approach can therefore complement a wide range of ecological approaches (e.g., field observations, experiments, community science, etc.). In all, our study suggests that ecological metrics like floral richness can be efficiently monitored by combining machine learning with easily accessible publicly curated datasets (e.g., Flickr, iNaturalist).

## Introduction

Meadows are ecologically important, and are valuable ecosystems found in the montane regions of the world. They are in the transitional zone between forest and alpine areas, and at the elevations that support forbs, sedges, and wildflowers. Functionally, montane meadows offer essential services such as water regulation, carbon sequestration, and erosion prevention (Yongheng 2007, Hopping et al. 2018). They additionally provide habitat for wildlife and are thus crucial for restoration and conservation efforts (Hik et al. 2001, Marini et al. 2007, Rui et al. 2015). Finally, their captivating floral displays also draw visitors and hold cultural significance, particularly for indigenous communities. However, alpine environments housing montane meadows are also uniquely susceptible to climate change (Ganjurjav et al. 2016, Steinbauer et al. 2018).

Montane meadows are particularly sensitive to changes in temperature and precipitation patterns because biological activity is generally limited by a short growing window that is driven by late snowmelt or cold temperatures (Cardinale et al. 2007), making them particularly sensitive warming climate (Inouye 2008). Floral richness is one measure that provides an important yardstick on the fitness of Alpine wildflowers, as meadow species are generally perennial (Roslin et al. 2021). Furthermore, flowers are good indicators for measuring the effects of climate change (Dreyer et al. 2006, Miller-Rushing et al. 2008, Gallagher et al. 2009, Rawal et al. 2015). Studying montane meadows provides insights into ecosystem dynamics, plant and animal adaptations, and climate change effects (Stöcklin et al. 2009, Jiang et al. 2018, Duan et al. 2021). Their sensitivity to changing climate patterns makes them valuable indicators of climate change impacts, with shifts in plant communities and downstream effects offering early warnings.

Phenological shifts in flowering due to climate change could have dramatic ecological impacts (Gottfried et al. 2012, Anthelme and Dangles 2012) as well as sociological effects. An extremely warm year might hasten flowering and make flowers susceptible to frost damage (Klein et al. 2018). On other hand, in a year with late snowmelt (i.e., after an extremely snowy winter or an unusually chilly spring), flowering might get delayed, resulting in less time for flowers to develop into fruits. Years with earlier flowering phenological could also result in a mismatch between pollinators and flower, if pollinators emerge after peak flowering or earlier before the onset of flowering (Inouye 2008, Ogilvie et al. 2017, Vorkauf et al. 2021). Furthermore, the experience of human visitors who visit to enjoy the wildflower displays could be at risk after these mismatches (Breckheimer et al. 2020, Hille Ris Lambers et al. 2021). Hence, evaluating patterns of floral richness spatiotemporally is important for understanding the factors driving meadow species reproductive life cycle (Inouye et al. 2003).

Traditionally, wildflower richness and flowering phenology has been studied with labor-intensive and time-consuming field observations. Detecting flowers from pictures taken in natural settings (such as alpine meadows) could provide a solution but is a complex task that recent developments in computer vision algorithms may aid in. Isolating flowers in a picture of a typical mountain meadow is challenging because of the diversity of meadow species, the wide range of floral structures, and because of the backgrounds in which the flowers exist (like leaves, trees, rocks etc.). Flowers are inherently complex because of the extraordinary diversity of floral morphologies resulting from eco-evolutionary dynamics (Moyroud and Glover 2017). However, machine learning (ML) vision-based algorithms show promise in detecting life stages or phenologies (e.g., identifying blooming), and in-depth evaluation of a single phenological stage for particular species has been successful (e.g. fruit maturity in passion fruit (Tu et al. 2018), strawberries (Chen et al. 2019) and tomatoes (Wan et al. 2018); ripeness in papayas (Santos Pereira et al. 2018), and flowering in cotton plants (Jiang et al. 2020)). ML approaches have also been used for classifying the progression of flowering in apples (Dias et al. 2018, Wang et al. 2021). These studies showed that it is possible to evaluate stages of a plant life event (for e.g., fruiting) from images with ML methods. However, these approaches have not been attempted in more species rich and complex systems like Alpine wildflower meadows, where issues related to occlusion, complex backgrounds and a high floral richness may complicate these approaches (Osherov and Lindenbaum 2017, Picek et al. 2022).

An evolving class of machine learning algorithms called deep neural networks (dNNs) have been particularly useful for detecting and segmenting multiple objects in an image. dNNs have found numerous applications in ecological conservation (Clark et al. 2023). For example, advanced dNN algorithms, such as convolutional neural networks (CNNs) have been deployed for image and audio recognition while recurrent neural networks (RNNs) have been used to analyze sequence data. Both contribute to biodiversity monitoring by identifying species from images, audio recordings and genetic sequences sometimes extracted from the environment (Aide et al. 2013, Khalighifar et al. 2021). These approaches also play a pivotal role in analyzing satellite imagery for land cover classification and detecting deforestation, aiding in habitat mapping and environmental monitoring (Chiang et al. 2020, Kattenborn et al. 2021). Thus, dNNs can be harnessed to predict the impact of climate change on ecosystems, enabling more accurate modeling of how changing conditions affect a wide variety of biodiversity and ecosystem services (Anderson 2018). These applications highlight the potential of deep learning to enhance conservation strategies by extracting insights from complex ecological data.

Here, we explore the use of dNNs to facilitate conservation planning by assisting in the species identification from multimodal data (images, in our case). Similar applications exist, ranging from tree crown detection to identifying cancerous cells (Abdolali et al. 2020, Hao et al. 2021). Like many of these applications, we employ dNN methods to perform object detection (for e.g., a flower) and instance segmentation (e.g., detect all the instances of a flower). Our use of dNNs for mapping flowers is also motivated by their ability to extract subtle features and learn from them efficiently (Dias et al. 2018). This means flowers can be individually tagged and segmented in a meadow image with dedicated dNNs. Thus, our work contributes to a growing integration of deep learning techniques into ecological research which can provide opportunities to address crucial conservation challenges.

Broadly, dNN approaches to identify objects (such as flowers) can be divided into one-stage and two-stage techniques. One-stage techniques perform object detection in a single pass by directly predicting object classes and bounding boxes, offering real-time processing but potentially sacrificing precision for small or overlapping objects. In contrast, two-stage techniques follow a multi-stage process involving region proposal and refinement steps, enabling more accurate localization and handling of overlapping objects at the cost of increased computational complexity and slower processing. A widely used one-stage algorithm called YOLO (Redmon et al. 2016) is a well-known object instance detection method, but has issues with objects with multiple sizes (pictures where objects are taken at different distances), and background noise. RetinaNet is a comparable one-stage algorithm that overcomes these limitations (Iglovikov and Shvets 2018). However, these approaches do not fully utilize certain attributes for classification, especially if objects are close together or small (Zhou et al. 2019). In such cases, a two-stage algorithm like Mask R-CNN (He et al. 2017) becomes advantageous. By integrating feature extraction, segmentation, and pixel-level classification, Mask R-CNN has been shown to achieve higher accuracy in various studies (Jia et al. 2020, Machefer et al. 2020, Hao et al. 2021).

In this paper, we therefore explore these three state-of-the-art dNN methods (YOLO, RetinaNet, and Mask R-CNN) for the detection and localization of Alpine wildflowers in complex settings – i.e., montane meadows. We define identification of a species by its flowering phenophase only i.e., the stage where the species flower(s) are visible. Specifically, we focus on two objectives; i) Evaluating the three dNN based techniques in their ability to detect flowering species in alpine meadows to derive diversity and abundance and ii) using the data to explore an ecological hypothesis on the relationship between elevation and floral richness to demonstrate the utility of this approach. Our expectation is that diversity declines with increasing elevation but in a non-linear way, and that ML approaches applied to pictures can capture this relationship.

## 2. Materials and Methods

### 2.1 Study Area

We conducted our study in the subalpine regions (upward of 1400 m) of the Mount Rainier National Park (MORA), which have an impressive wildflower display that is one of the main summertime visitor attractions. Meadows are snow covered most of the year, even late into July, with peak-flowering that lasts between 2 and 3 weeks between late July and early September – depending on snowmelt dates (Theobald et al. 2017). We collected data along an existing network of trails called the Reflections Lakes Trail, which has been a site for focused meadow phenology observations (Hille Ris Lambers et al. 2021, Manzanedo et al. 2022). The trail is on the south side of MORA and covers an elevational gradient of 1400 m to 1950 m (Fig. 1).

**Figure 1:**
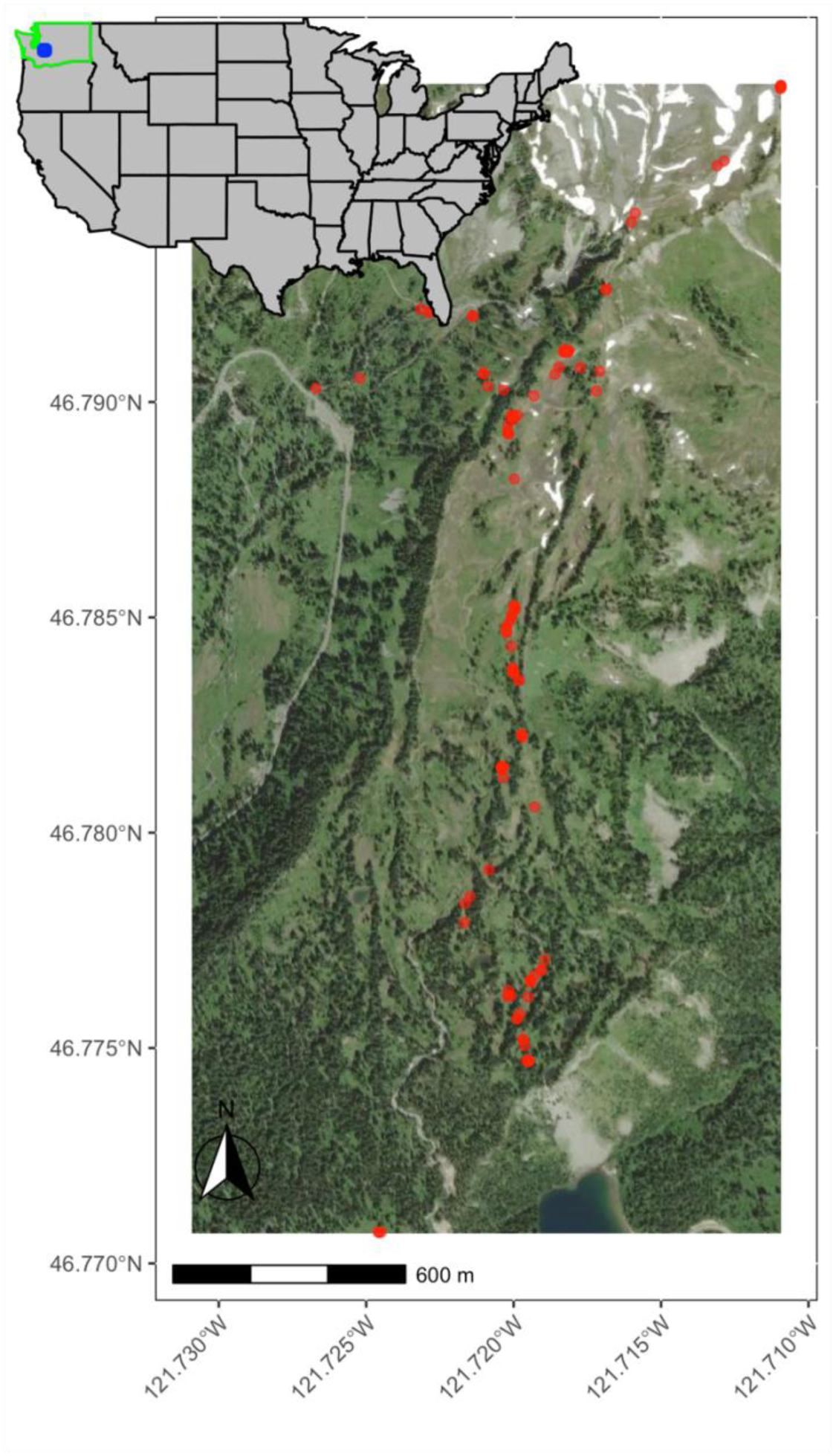
Study area at Mt Rainier National Park, with the red dots indicating the sites along the Reflections Lakes trail at which the pictures were taken for the study (n=27). The inset shows the location (in blue) of the study area in the context of Washington State, USA.

### 2.2 Data

Volunteers participating in this study collected camera images using an iPhone camera (version 11 Pro) during the flowering season of 2020 and 2021. Sixteen of the 42 species that commonly occur at the study site were targeted in this study (see Table 1). The subset was chosen based on species list in use by the community science program MeadoWatch (Manzanedo et al. 2022). Photos were taken from the trail overlooking the meadows. The curated dataset contained 1545 high resolution photos: 768 x 1024-pixel (landscape) and 1024 x 768-pixel (portrait). The portrait and landscape images were then tiled for easier annotation after converting them to a common dimension of 1024 pixel x 1024 pixel. Padding was done so that it was easier to tile the images in equal sizes. For e.g., tiling into 256 x 256 of a 1024 x 1024 format image results in 8 images. We partitioned the data into training, test, and validation sets. Specifically, 70% of the data was allocated to the training set, while the remaining 30% was evenly distributed between the test and validation sets. In total, we used 1221 images for training and 324 images for validation and testing.

**Table 1:**
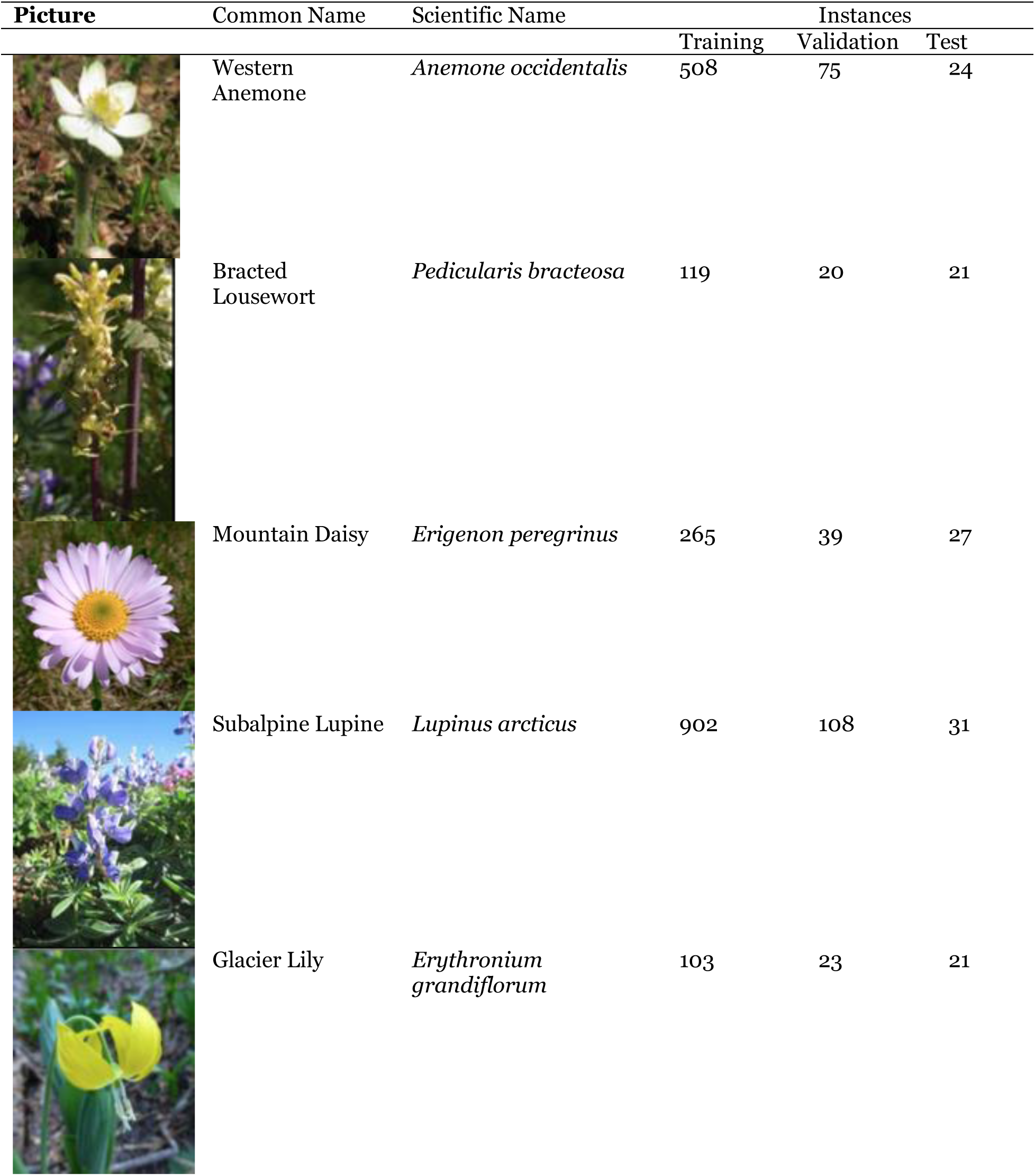
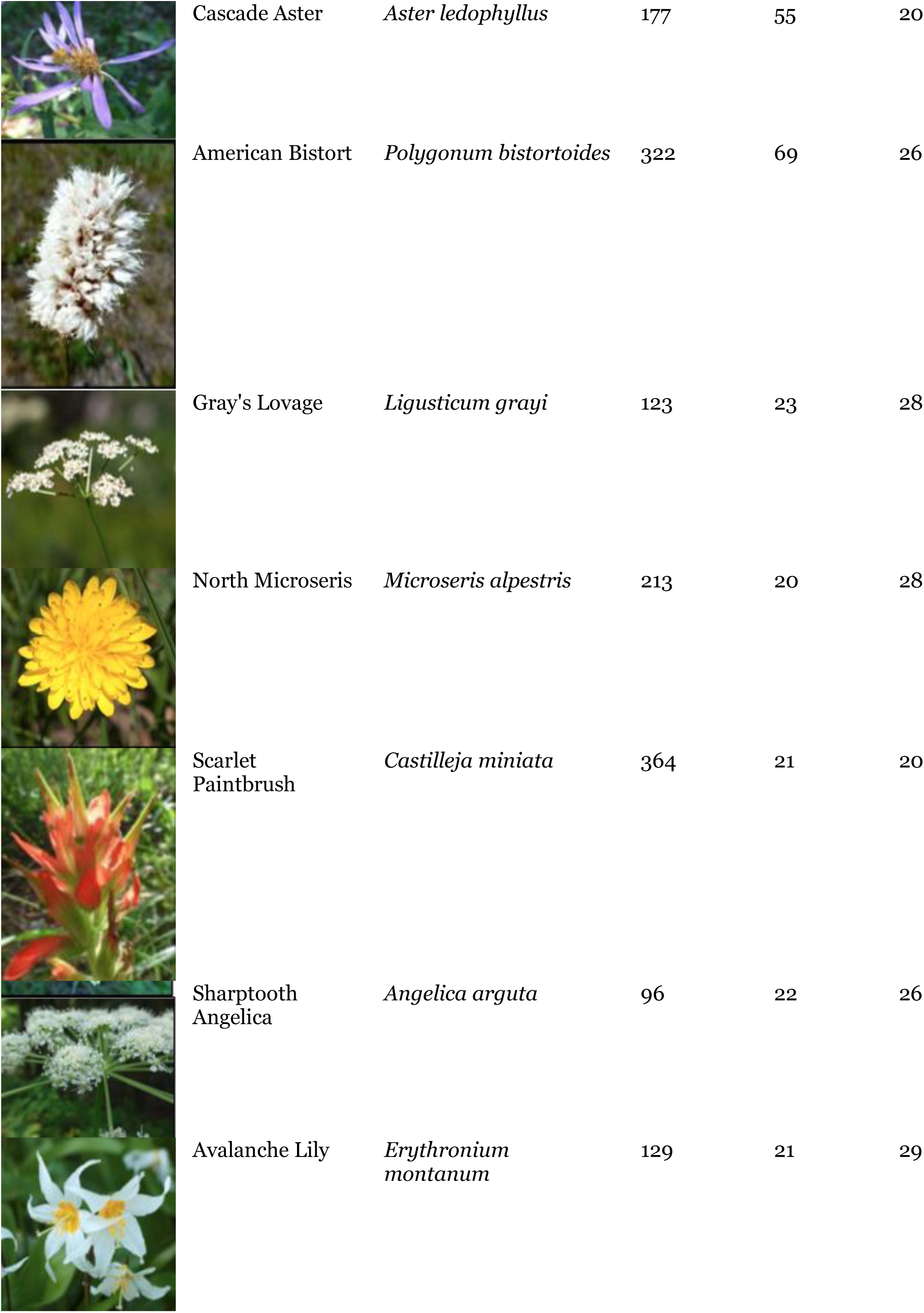
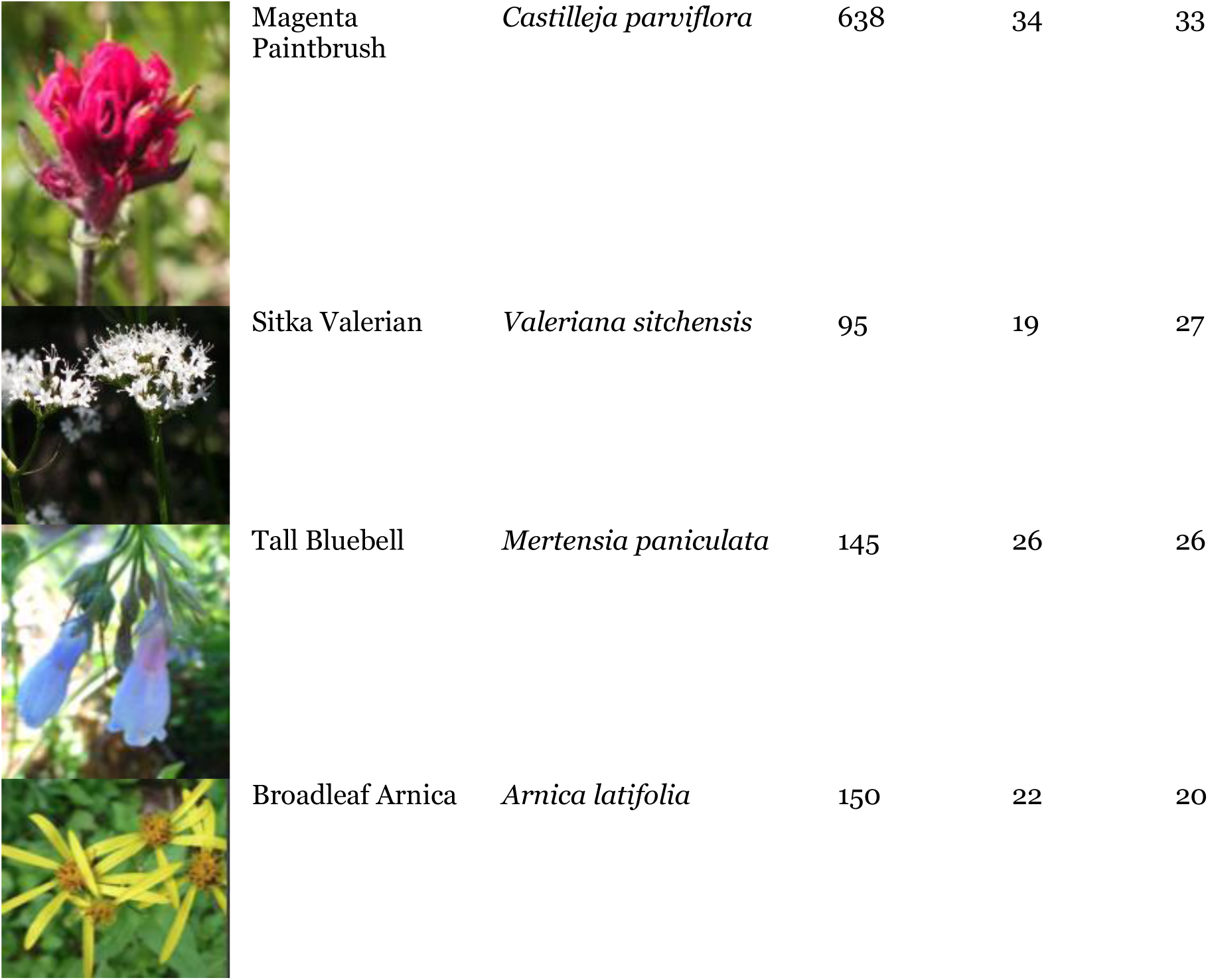
The training dataset comprises a total of 1221 images having 4349 instances of flowers annotated. The validation dataset had a total of 166 images having 597 instances of flowers annotated, and the test dataset consisted of a total of 158 images having 407 instances of flowers annotated. All the images were 256 by 256 pixels and collectively represented 16 wildflower species.

For our training dataset, we annotated each individual image, identifying the number and identity of flowers in each image. We used the LabelMe software to perform the annotation (Russell et al. 2008), which was then outputted in a JSON (JavaScript Object Notation) file format. These annotations included multiple labels (flowers) with their annotation mask (see Fig. 2 for sample annotations). For the application of the Mask R-CNN and the RetinaNet dNN algorithms, the annotation outputs were in JSON format, which was converted to a COCO format by the training framework. Because the YOLO algorithms require a specific format (the YOLO format), we converted the LabelMe annotations to this format prior to analysis.

**Figure 2:**
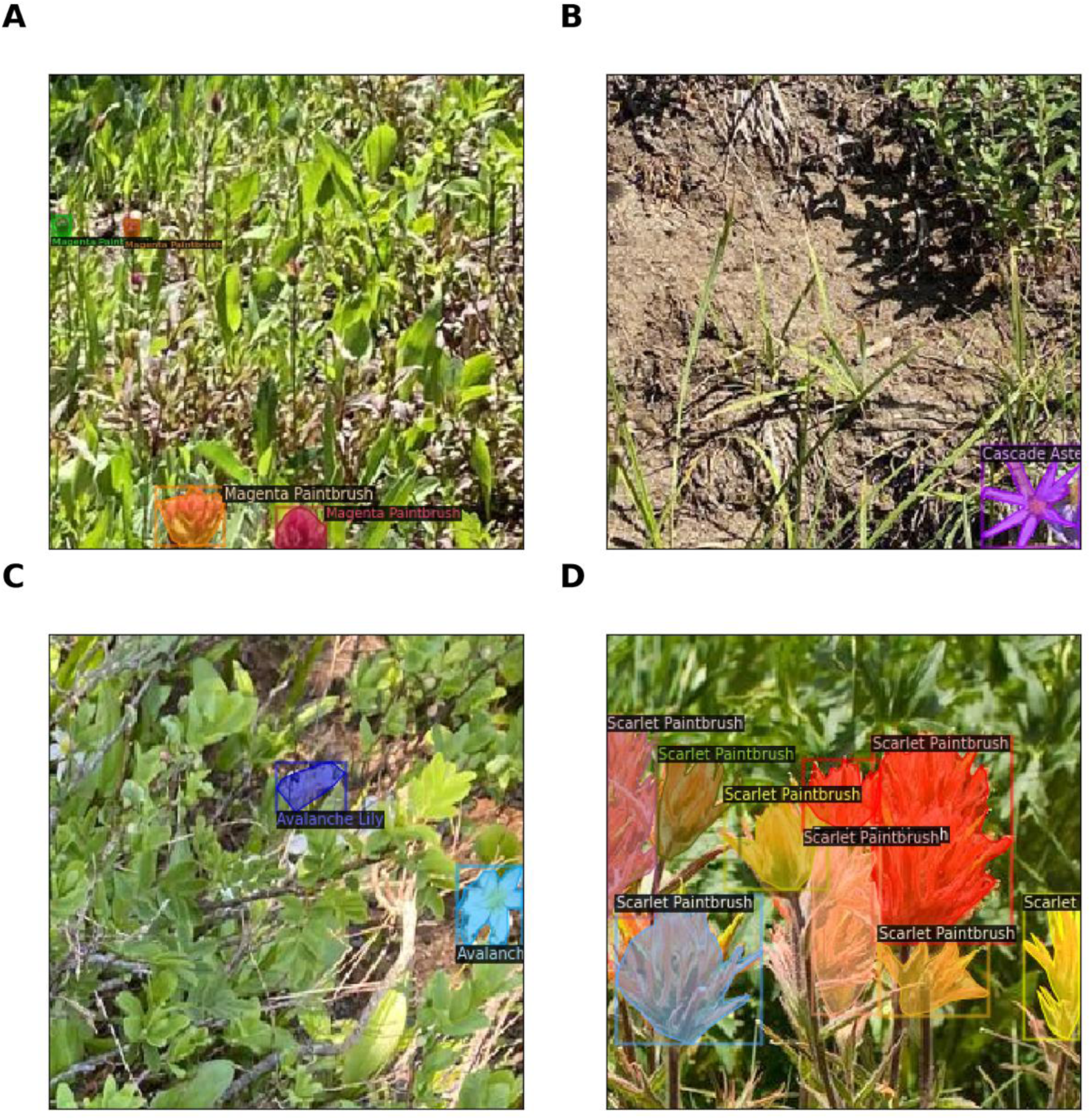
Example annotated images from the training dataset. (A) Magenta Paintbrush (*Castilleja parviflora*) flowers (B) A flowering Cascade Aster (*Aster ledophyllus*) (C) Avalanche lilies (*Erythronium montanum*); particularly found earlier in the season (D) Cluster of Scarlet Paintbrush (*Castilleja miniata*) in full bloom. Collectively, these images illustrate the complexity of the machine learning task - with different abundances, photo angles, and backgrounds.

### 2.3 Machine Learning Methods

We explored how three dNN algorithms performed in detection and segmentation: YOLO, RetinaNet and Mask R-CNN. These three methods are inherently able to extract features and learn from them efficiently (Dias et al. 2018). In our study, we utilized YOLOv5 (here onwards referred to as YOLO), which solely supports bounding boxes. However, the latest version is YOLOv8 that also supports generation of pixel level mask (segmentation). YOLO and RetinaNet support bounding box detection only, whereas Mask R-CNN and YOLO give the mask of the detected object in addition to the bounding box. We briefly describe the three methods in Appendix A. We used the Detectron2 framework (an implementation of the PyTorch deep learning library) from Facebook Artificial Intelligence Research (FAIR) to train deep neural network models for flower detection for RetinaNet and Mask R-CNN. The Detectron2 has an out-of-the-box implementation of many of the latest CNN algorithms that perform detection and segmentation (Wu et al. 2019). Detectron2 allows writing reproducible code with a standardized way of dealing with input data and annotations. Detectron2 also provides boiler plate code for dataset handling, augmentations, training, visualization, and performance metrics. Augmentation methods enhance the versatility of the model. We applied two augmentation techniques during training: random image flipping and rotation (Xu et al. 2023). All three of these methods i.e., YOLO, RetinaNet, and Mask R-CNN apply bounding box detection (i.e., localize the object), but Mask R-CNN and the recent version of YOLO (v8), in addition, gives pixel level masks of the class. In addition, Mask R-CNN supports pluggable feature extraction network backbone like U-Net (Ronneberger et al. 2015) that could cover a wide range of domains. The development, training and inference of the models were performed on Azure Cloud on a server having the configuration Standard NC12_Promo (12 vCPUs, 112 GiB memory with 2 NVIDIA Tesla K80 GPUs).

### 2.4 Hyperparameter tuning

We performed hyperparameter tuning using the validation data composed of 597 instances (compared with 4349 instances used in training). Hyperparameter fine-tuning of all three models was achieved with training for 1500 iterations on Google Colab’s K80 GPU. Anchors are predefined with aspect ratio and scale to establish the location and size of the bounding box for an object. We used different anchor sizes (Kalantar et al., 2020) for the three algorithms. We kept the learning rate at the default value of 0.00025 with a batch size of 128. We also kept the aspect ratio of 1:1 and scale to double with a total of 5 increments.

### 2.5 Evaluation Metrics

We used detection evaluation metrics as defined by the COCO dataset (Lin et al., 2014). Intersection over Union (IoU) is a metric that evaluates how the ground truth and predicted detections overlap. IoU of 1 means an ideal detection whereby there is an exact overlap whereas IoU of zero means that there is no overlap. Average Precision (AP) indicates how a detection method performed at a particular IoU. Here, we report AP at 0.5, 0.75 and 1 (as in Table 2). AP is calculated by taking average of AP of ten IoU thresholds (t= [0.5,0.55,0.60,0.65,0.70,0.75,0.80,.0.85,90,.95] and when averaged across all categories the metric provides us mean average precision (mAP). True positive (TP) would mean IoU > = Threshold, and false positive (FP) is IoU < Threshold. We further inspect the Precision (TP/TP+FP) and Recall (TP/TP+FN) to evaluate frequency of false detections or missed ones; note that higher mAP indicates better detection performance. Average Precision (AP) and Average Recall (AR) is further used to calculate the F-1 score which helps with interpretability of the techniques. Mask R-CNN identifies the anchor boxes based on the IoU, which is a critical step in the first part of the network (RPN).

**Table 2:**
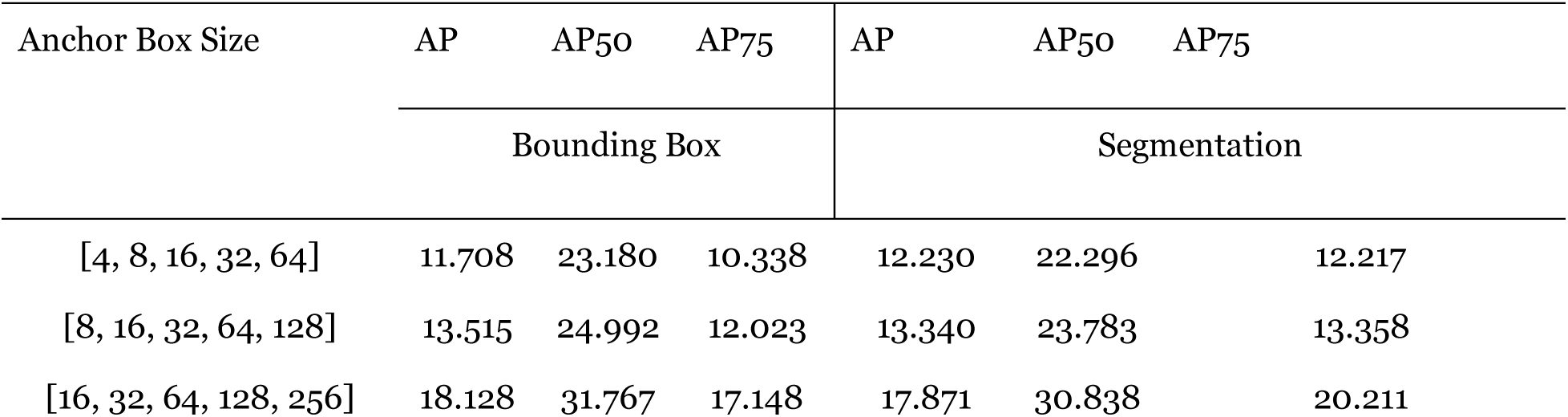
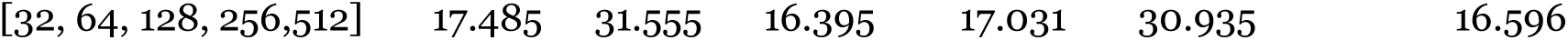
Results of anchor box variation with Mask R-CNN.

For each photograph, we computed two floral indices: species diversity, as quantified by Shannon’s diversity index, and species richness. The Shannon diversity index was determined using the formula 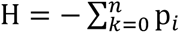 ln p_*i*_, where p_*i*_ represents the proportion of a specific flowering species in the photograph (Shannon 1948). Species richness was determined as the count of distinct flowering species within a given photograph.

## 3. Results

### 3.1 Hyperparameter tuning results

We found that maintaining anchor box variations that are multiple of 16 i.e., [16,32,64,128,256] yielded better accuracy (Table 2). The results were consistent with YOLO and RetinaNet (Appendix C).

### 3.2 Performance and training time of the flower detection methods

We tested the three dNNs algorithms using over 316 images with an average number of 40 labels per flower category. We used two augmentations in training: Resize Shortest Edge and Random Flip. Each of the augmentation methods were trained for 10000 epochs. The time needed for training differed between the three algorithms, ranging from 3 hours for YOLO to 12 hours in Mask R-CNN.

Table 3 shows the mAPs and different IoU thresholds for all the methods. In addition, Mask R-CNN provides a classification metric by identified object size as well. In our case, for small, medium, and large flowers, the mAP was 29.27, 42.76 and 48.80, respectively. Bounding box detection applies to all the methods, but segmentation (drawing of masks) only applies to Mask R-CNN, so segmentation scores are only reported for Mask R-CNN.

**Table 3:** Evaluation results for the BBox and Segmentation across the three algorithms. Mean Average precision (mAP) for the three methods. All three algorithms were trained for 10000 iterations (epochs).

| Method | BBox |  |  | Seg |  |  | F-1 Score | Training time (hours) |
| --- | --- | --- | --- | --- | --- | --- | --- | --- |
|  | mAP | mAP50 | mAP75 | mAP | mAP50 | mAP75 |  |  |
| Mask R-CNN | 45.80 | 67.24 | 52.10 | 31.32 | 59.05 | 30.78 | 0.20 | 12 |
| RetinaNet | 23.59 | 51.28 | 18.63 | N/A | N/A | N/A | 0.12 | 8 |
| YOLO | 32.66 | 60.68 | 31.91 | N/A | N/A | N/A | 0.23 | 3 |

All three methods were able to isolate the flowers within the visual complexity of a meadow. Table 4 gives the breakdown of mAP by species for all the three methods. On average, Mask R-CNN performs better for all species, but we note differences in identifying individual species across the methods. For example, Scarlet Paintbrush, Tall Bluebell and North Microseris are detected more accurately by YOLO than by Mask R-CNN or RetinaNet. Regardless of the species, they are detected with higher accuracy if they are towards the foreground of the image and if the image is in focus - see for example Bracted Lousewort being isolated by Mask R-CNN while two species in the horizon are missed (Fig. 3A, B). Similarly, occluded flowers are being missed for example some of the Mountain Daisy flowers that are behind the stalks of other species (Fig. 3C, D). The sample prediction results for YOLO and RetinaNet are in Appendix B.

**Figure 3:**
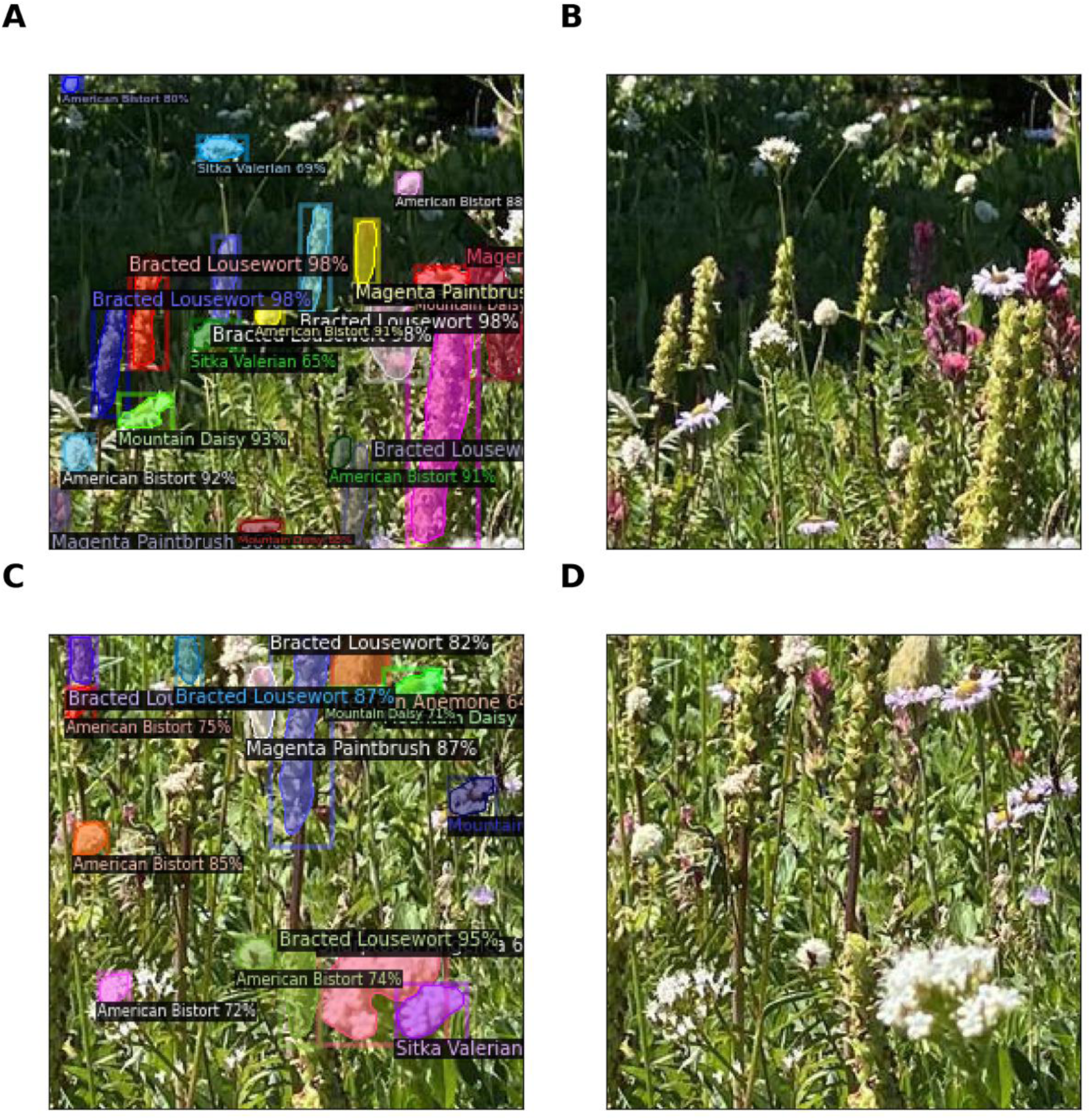
Predictions from the Mask R-CNN method showing the fluency of the model; the percent next to the predicted class represents model confidence. (A and B) Meadow in peak flowering with predictions (B) corresponding original tile. (C and D) Another meadow in peak flowering with predictions (D) corresponding to the original tile.

**Table 4:** Per category BBox evaluation (using mAP) results across all three methods.

| Flower | AP50 |  |  |
| --- | --- | --- | --- |
|  | Mask R-CNN | RetinaNet | YOLO |
| *Western Anemone* | 58.45 | 35.22 | 27.90 |
| *American Bistort* | 42.79 | 23.55 | 56.80 |
| *Avalanche Lily* | 40.30 | 18.74 | 52.2 |
| *Bracted Lousewort* | 58.05 | 18.28 | 4.42 |
| *Gray's Lovage* | 36.02 | 20.07 | 43.90 |
| *Magenta Paintbrush* | 42.01 | 29.98 | 27.80 |
| *Mountain Daisy* | 51.42 | 23.31 | 67.70 |
| *North Microseris* | 68.22 | 39.10 | 77.80 |
| *Sitka Valerian* | 45.43 | 21.31 | 10.40 |
| *Subalpine Lupine* | 38.36 | 24.67 | 39.20 |
| *Scarlet Paintbrush* | 59.91 | 33.00 | 74.60 |
| *Tall Bluebell* | 46.04 | 14.45 | 83.00 |
| *Glacier Lily* | 29.93 | 11.56 | 14.70 |
| *Sharptooth Angelica* | 24.34 | 08.11 | 48.30 |
| *Broadleaf Arnica* | 61.09 | 21.80 | 41.50 |
| *Cascade Aster* | 26.83 | 34.34 | 07.62 |

### 3.3 Richness and abundance of flowering species

We found that floral species richness and abundance per photo varies along the elevational gradient. Specifically, we see lower floral richness at lower and higher elevations, and increased richness along mid elevations i.e., 1700-1800 m (Mask R-CNN shown in Fig. 4A). YOLO aligns with Mask R-CNN on elevation and richness relationship (Fig. 4A, C and Appendix D), but RetinaNet does not suggest any association (Fig 4E), We also see increased density of flowers at the mid-elevation compared to lower and higher elevations (Fig. 4B). RetinaNet showing elevated density of flowers compared to Mask R-CNN and YOLO (Fig. 4B, D, F). Floral abundance in pictures was also found to vary over time, even over very short periods with similar overestimation by RetinaNet (Fig. 5).

**Figure 4:**
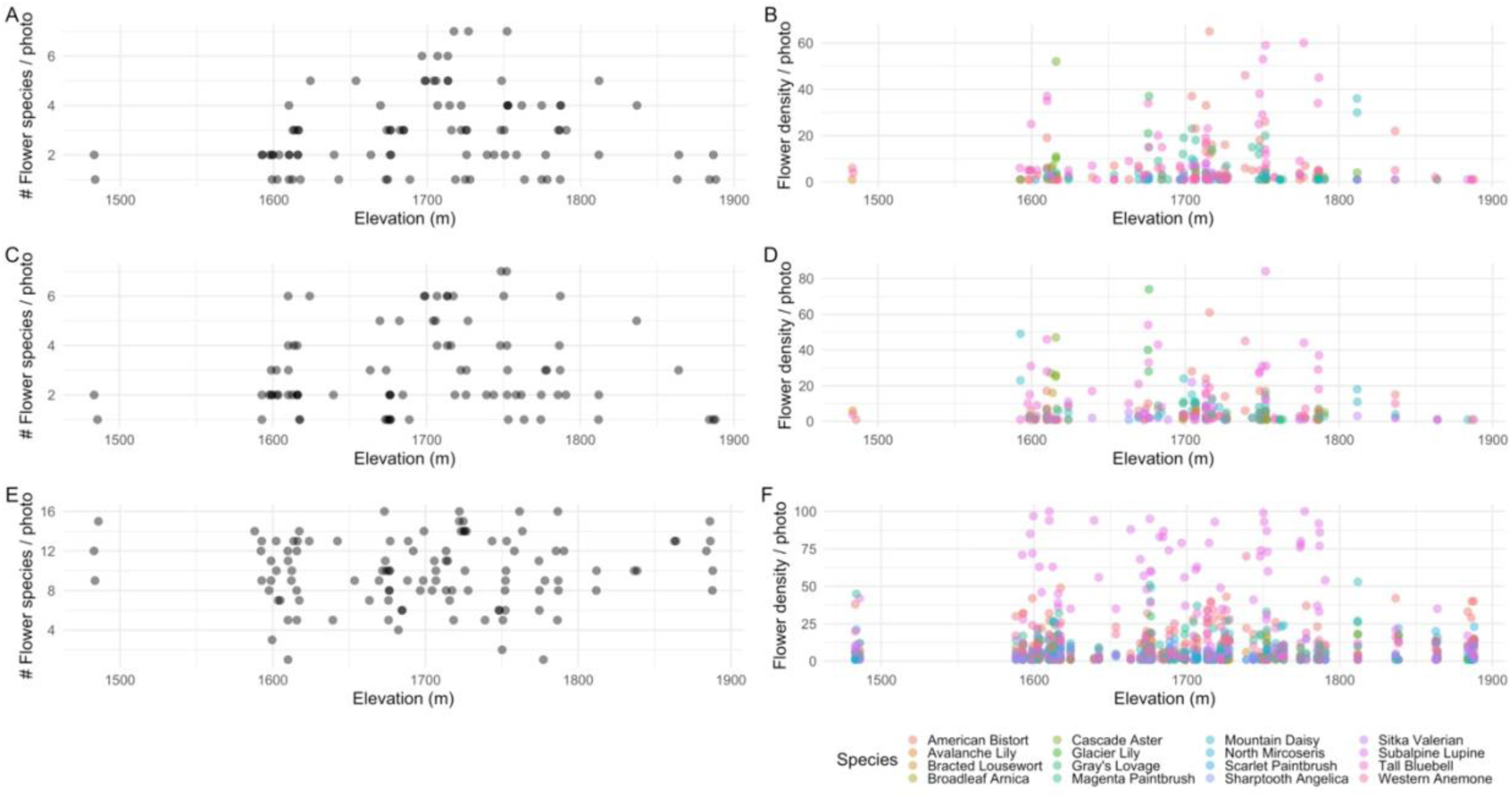
Detected species from all the collections using images collected in August 2020; predictions are from Mask R-CNN, YOLO and RetinaNet methods. Number of unique flower species detected per photo (A) by Mask R-CNN, (C) by YOLO, and by RetinaNet (E). Number of occurrences of each flower species per picture (B) by Mask R-CNN, (D) by YOLO, and (E) by RetinaNet.

**Figure 5:**
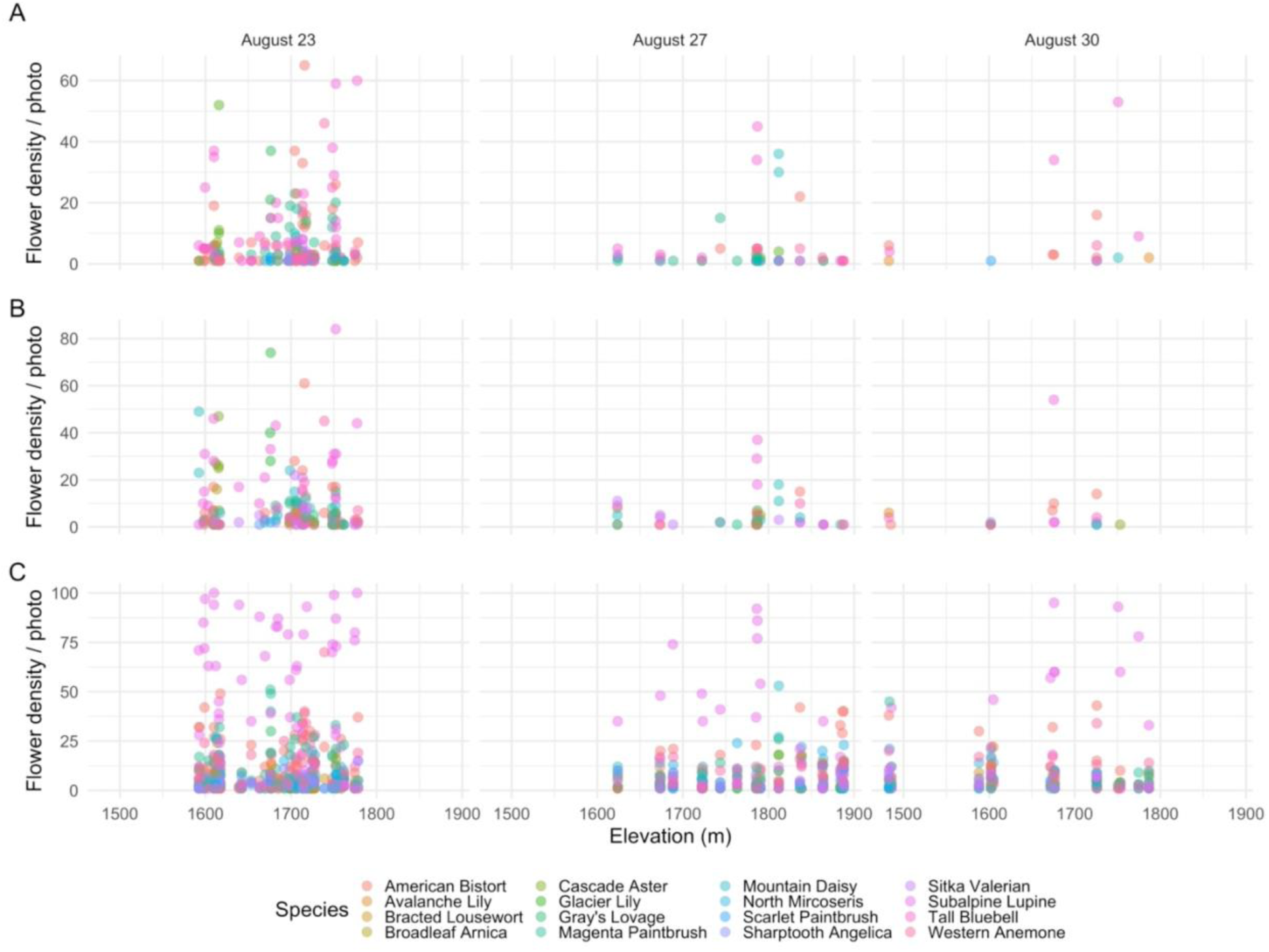
Floral density of species by picture for all the collection dates in August 2020 from all three methods, (A) by Mask R-CNN, (B) YOLO (C) RetinaNet.

We found a significant and positive correlation between diversity, determined through annotations, and diversity as assessed by predictions from dNN methods using a separate labeled dataset (correlation coefficient R = 0.62 for Mask R-CNN, R = 0.52 for YOLO, and R = 0.1 for RetinaNet, all with a p-value of less than 0.001; refer to Fig. 6A). Among the dNN methods, this correlation in terms of species richness was evident for Mask R-CNN and YOLO (correlation coefficient R = 0.36 for Mask R-CNN, and R = 0.42 for YOLO, both with a p-value of less than 0.001; see Fig. 6B), but such a correlation was not observed for RetinaNet. Moreover, there existed a correlation between Shannon diversity and species richness of flowers when comparing Mask R-CNN and YOLO (correlation coefficient R = 0.51 for Shannon diversity and R = 0.8 for species richness, both with a p-value of less than 0.001; refer to Fig. 6C and Fig. 6D, respectively).

**Figure 6:**
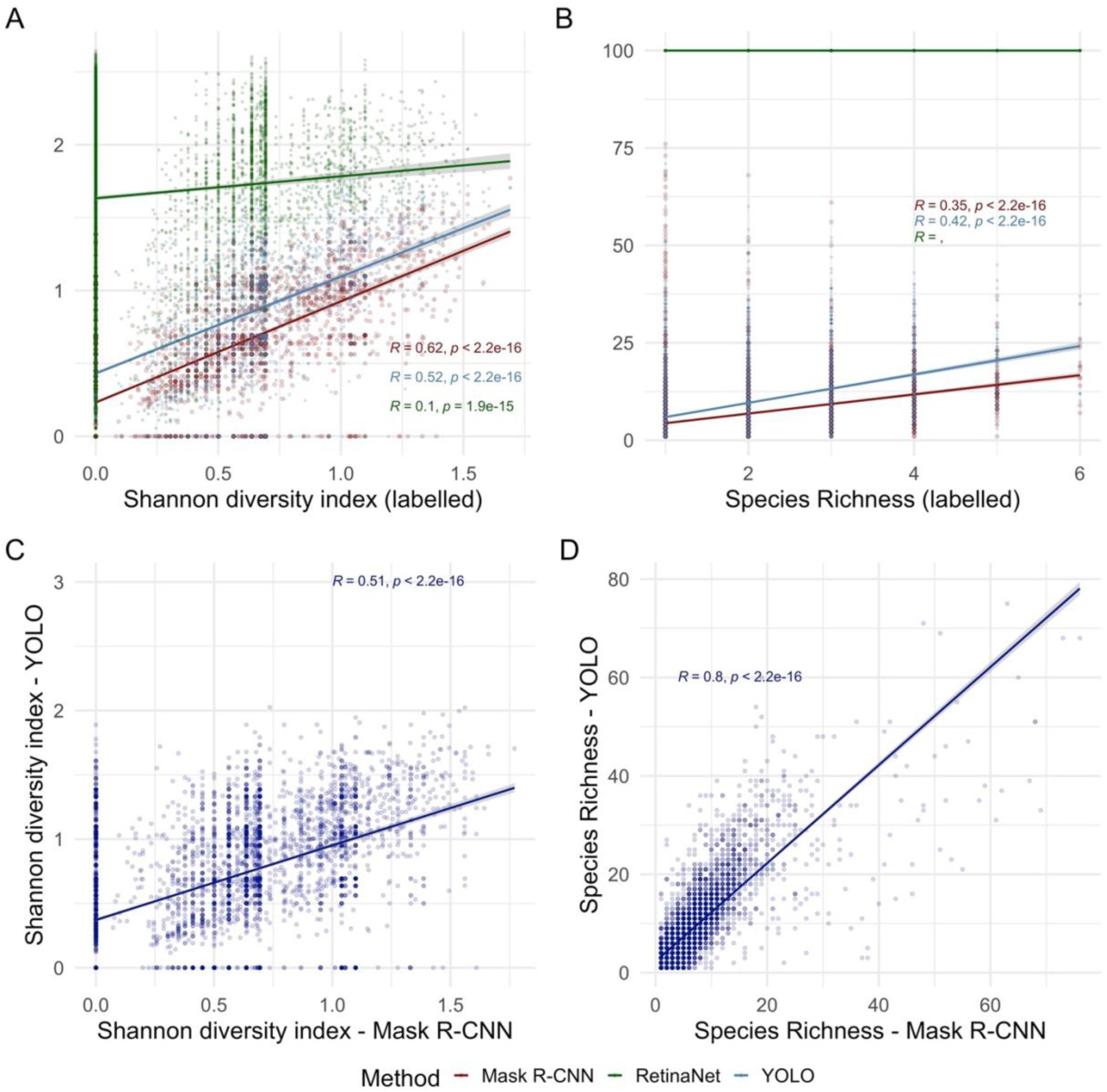
Comparison of diversity and richness using a labelled hold-out dataset across all three methods, (A) Diversity, (B) Richness, (C) Diversity between YOLO and Mask R-CNN, and (D) Richness between YOLO and Mask R-CNN. Inset showing Pearson’s correlation coefficient, and the corresponding p-value.

## 4. Discussion

We demonstrate that deep learning techniques can detect, identify, and count Alpine wildflowers in photographs, and thereby provide detailed information on flowering occurrences in complex mountain meadow systems. In general, Mask R-CNN demonstrates superior performance; however, YOLO and RetinaNet exhibit better results for specific flower cases. Furthermore, we show that these approaches verify known ecological patterns – that species flowering varies temporally and by elevation at our study site (Kudo and Suzuki 1999, Prevéy 2020). Our results show that with timestamped, geo-coded photographs, deep learning techniques can identify and count flowers, potentially facilitating detailed analyses on flowering richness as well as floral abundance, both key ecological metrics relevant to Alpine wildflower fitness and pollinator performance.

The relative performance of the three dNN methods we tested depended on task-specific requirements, performance-tradeoffs, and the necessity for detailed instance segmentation. For object detection and instance segmentation, YOLO, and Mask R-CNN stand out as equally implementable methodologies, with variation in performance. YOLO’s real-time efficiency derives from its single-pass approach, making it well-suited for rapid detection tasks; however, challenges with this approach arise in accurately identifying small and densely overlapping objects. RetinaNet addresses scale variability by incorporating anchor boxes and feature pyramid networks, finding equilibrium between speed and precision, though it might not excel in intricate instance segmentation. In contrast, Mask R-CNN’s offers unparalleled pixel-level accuracy for object separation but demands greater computational resources and time. We discuss these points in more detail below.

### 4.1 Differences in performance and methodologies for identifying flowers in camera images

In our application, the two-stage detector (Mask-R-CNN) was more accurate but slower in its predictions than the one stage detectors (YOLO, RetinaNet). This is consistent with other studies using this approach to count and identify individual flowers of a single species (Yu et al. 2019, Machefer et al. 2020, Hao et al. 2021). Single stage detectors perform classification and regression together on the predicted anchor boxes. The trade-off is that adding an additional stage incurs computation time (Suh et al. 2021). The YOLO algorithm which is a single stage detector is able to accurately draw boxes around the flowers as well as classify them correctly more speedily than Mask R-CNN (similar findings are presented in ((Prasetyo et al. 2020) and (Sumit et al. 2020)). However, it is crucial to emphasize the significance of annotations. Predictions made within images might inadvertently miss numerous instances of flowers, a situation expected due to the inherent complexity in annotating such occurrences during the training phase.

The images acquired for our study were in two dimensions (1024 x 768 and 768 x 1024). However, we found that breaking down these extensive format images into 256 x 256-pixel images resulted in the lowest classification error. The 256×256 images seem to offer sufficient dimensions for detecting objects like flowers of similar size within the dataset. We observed that dealing with larger format image samples demanded more computational resources. While analyzing the metrics, it is important to consider that larger images lead to higher occurrences of box and objectness errors. This is because sub-images are extracted from the same larger image, and if the sub-image is larger, it will contain more boxes within it (Safaldin et al. 2023). As these errors are aggregated for each potential box, the cumulative box and objectness error tends to be greater for larger images.

While our deep learning study was done with a minimal dataset, the initial results are promising. Augmenting the dataset through more annotation is apparent, but the resource-intensive nature of this step needs to be weighed against computation and resource availability. Moreover, exploring algorithms designed for efficacy under weak supervision (heuristically labeled data) could be a potential avenue for future enhancement (Phan et al. 2023).

### 4.2 Flower characteristics, species abundance and implications for detection

Several factors led to variation in our ability to classify and detect flowers. Most wildflower meadows (and thus pictures) include large variations in the abundance of species, like Subalpine Lupine, which tends to be more abundant than other flowers. This results in Subalpine Lupine being detected more than other species like Mountain Daisy which appears more sparsely, a finding that reflects real differences in floral abundance. However, the algorithm also makes mistakes in species identity, and it is likely that the similar characteristics of some flowers are responsible. For example, some meadow wildflowers have an elongated floral structure (e.g., Bracted Lousewort and Subalpine Lupine), while others are more circular in nature (e.g., Mountain Daisy and Cascade Aster). Misclassification tended to occur when structures and color were similar (e.g., Sharptooth Angelica and Gray’s Lovage). Unlike in other studies, in which data enhancement helped prevent overfitting and improved accuracy (Tian et al. 2020), we did not see improvement in accuracy with image augmentations. We also see the shortcoming where occluded flowers were not being isolated (Fig. 3A, C).

Despite these challenges, we were encouraged by the accuracy we achieved in this study as these pictures are inherently complex and therefore challenging for classifiers (Fig. 6). Some challenges include inconsistent light levels (e.g., because of slope and aspect) and heterogenous backgrounds (e.g., rocks, lots of plants, etc.). Additionally, labeling errors, namely flowers that are missed while annotating could contribute to decreased accuracy. For example, the YOLO method penalizes incorrectly drawn boxes - if a flower is not labeled in the training set, YOLO would penalize the model for drawing a box around the flower despite it being correct. These challenges are all ones that could be addressed in studies that standardize photo acquisition and automation and increase annotation effort, both factors feasible for many ecological studies.

There also is likely to be sampling bias in this study. In the iconic wildflower meadows of Mt. Rainier National Park, the chances of picking a location and photographing it is almost certainly positively correlated with the probability of a flower existing in that place. That means that our pictures are disproportionately likely to contain flowers, potentially predisposing our algorithm to bias toward flowers. Another limitation of our study is that it did not capture the full flowering season and does not quantify the flowers by location (e.g., elevation). Fortunately, these issues are easily addressed with a planned study, for example with images are captured from intentionally placed PhenoCams along an elevational gradient.

### 4.3 Applicability to ecological research

We believe that combining image analysis with automated floral detection can provide important information on metrics of ecological interest, which can either complement or in some cases even replace more traditional ecological sampling. For example, our analyses show that floral richness and abundance are linked to elevation in a non-linear way (Fig. 4A, B), similar to patterns shown by other studies (Fierer et al. 2011, Guo et al. 2013, Antonelli et al. 2018, Sponsler et al. 2022). Floral richness is driven both by plant diversity (which is influenced by many biotic and abiotic factors), as well as by the probability that species are flowering (Fabbro and Körner 2004, Sargent and Ackerly 2008, Theobald et al. 2017) – an important indicator of plant fitness resources for pollinators. Thus, this approach could be used to provide information on diversity patterns across gradients, phenological patterns of flowering relative to climate, and floral resources for pollinators.

For abundant species with recognizable flowers, these approaches may also allow for the detection of distributional shifts, if conducted annually. For example, climate change might alter the distribution of specific wildflowers found in montane environments; possibly as the result of elevational shifts (Gottfried et al. 2012, Steinbauer et al. 2018). Similarly, a rapid invasion of a non-native flowering species can be detected using this approach. The combination of the floral detection approaches introduced here with drone imagery could allow for assessment of such shifts over large spatial areas, or in remote areas.

However, there are some caveats to using flowers in pictures to quantify richness and / or abundance. For one, floral richness is not total species richness, as many perennial plants do not flower every year or in every location. Moreover, different species produce differing number of flowers per individual, meaning that a higher number of flowers need not mean a greater density of individuals. A second important caveat in our measures of floral abundance is that fields of view vary from picture to picture, which means the total area available to flowers (and thus flowering abundance) will vary even if underlying patterns detected by more traditional ecological approaches (e.g., m^2^ quadrats) do not.

In total, however, we believe the promise of these approaches for generating large volumes of data with less manpower (as described in (Wilson et al. 2017)) outweigh these issues. Moreover, these challenges can be overcome with large volumes of pictures / data or by employing fixed cameras (e.g., webcams, PhenoCams) with common fields of view. For example, if pictures are taken automatically via PhenoCams or TrailCams, these fixed cameras allow for pictures to be taken in the same place repeatedly, which saves time and gives an advantage over taking them oneself as one gets to control when and where the pictures are to be taken. Then, differences in floral richness would not be due to differences in field of view (the way the picture was taken). Regardless, we feel that floral richness derived from photos is still of interest as it gives a quick proxy of site-specific flowering patterns.

We also believe our approach can also be used to address questions about phenology that otherwise would not be possible with traditional ecological sampling. As photos are engrained with date/time and location information, we could also have linked these measures to time to provide information on phenology (as in (Breckheimer et al. 2020)). In this case, crowd sourced pictures (e.g., Flickr, iNaturalist, Instagram) increased the data volume (# of observations) and allowed for quantification of phenology in previous years (i.e., going back in time). This and similar approaches, however, require specific statistical approaches. For example, crowd sourced pictures (e.g., from Flickr) are not taken according to a specific experimental design, which means using these pictures could result in an unbalanced representation – e.g., spatial or temporal gaps in distributions (e.g., Mt. Rainier meadow pictures predominantly may be from around visitor centers, weather permitting times and might have weekend effects). However, one can deal with skewness introduced by source images by statistically controlling for any spatiotemporal bias in pictures taken (Breckheimer et al. 2020).

Finally, we believe that the floral detection combined with ML approaches could allow for quick assessment of ecological patterns, which could be important in a changing world. For example, it might be valuable to know about the distribution of a rare species threatened by climate change and / or the co-occurrence of wildflowers that compete for pollinators whose phenology shift at different rates or due to climate driven changes in species composition (Taheri et al. 2021). Documenting species presence-absence using any of our methods as applied to existing (crowd-sourced) pictures could address questions such as these and provide a preliminary assessment of patterns before designing new (and costly) experiments (Elliott and Davies 2019).

### 4.4 Future directions

Although our flower detection algorithm performed well, there are likely improvements that could be made to detect a greater number of wildflowers and better distinguish between similar looking flowers. For example, we have not looked at the fluorescence signal but hypothesize that shape and color are being learnt during feature extraction, and as future directions, incorporation of additional information (e.g., thermal, or infrared bands) might help in the presented algorithms.

Embedding the trained models in community science apps (Mobile Applications) could also be one of the end uses of these algorithms. In other domains, apps have increased community involvement, e.g., the bird identification app, Merlin (Nugent 2018) and the plant identification functionality of iNaturalist. Use in apps both provides the public with a way to identify flowers and a way to gather data.

Combining photo data with drone and remotely sensed products could potentially offer opportunities to measure things in remote areas and on larger spatial scales. Improvement of algorithms that can use additional bands (produced in hyperspectral or multispectral imagery) could be promising and might detect more kinds of flowers and distinguish between flowers that are more similar (Wieme et al. 2022).

## 5. Conclusions

We explored three deep learning techniques to detect and count flowering plants in alpine meadows. The two-stage detector we used (Mask R-CNN) performed better than the single-stage detectors (YOLO and RetinaNet), with the first stage cropping flowers and filtering out the background; and the second stage classifying and detecting the flowers. However, all techniques gave us detailed information on flowering occurrences in meadows, allowing us to calculate floral richness as well as flower abundance. These approaches were also able to capture known ecological patterns; namely that flowering varies temporally (i.e., phenology) and by elevation. While our trained models exhibit promise for application in areas dominated by our focal species from Mt. Rainier National Park, caution should be exercised considering potential variations in local ecological dynamics. Looking forward, further investigations into the suitability and transferability of these models to subalpine regions across the broader Pacific Northwest (such as Washington, Oregon, and British Columbia) will be essential for robust and regionally informed study efforts. In the future, including a larger catalog of species for training could allow for coverage from other areas, for example Rocky Mountain meadows or Swiss Alpine meadows. More generally, these approaches could aid in climate change studies, because they can be used in a non-invasive, low-investment way to rapidly get information on spatiotemporal flowering patterns.

## Acknowledgements

We want to acknowledge support of the University of Washington eScience Institute and Hille Ris Lambers lab in developing the ideas and supporting discussions. Authors want to acknowledge assistance from John H., Sapti S., Tommy L., and Richard L. Cristea was supported by the NSF award OAC-2117834 and the University of Washington eScience Institute. We also acknowledge support from Microsoft AI for Earth and Microsoft Azure collegial support. This study was carried out on the ancestral lands of the people of the Nisqually, Puyallup, Squaxin Island, Muckleshoot, Yakama, and Cowlitz Nations.

## Declaration of Competing Interest

The authors declare that they have no known competing financial interests or personal relationships that could have appeared to influence the work reported in this paper.

## Authors’ contributions

A.J. and J.H.R.L. conceived the ideas; A.J., J.H.R.L., N.C., A.T. designed the methodology; A.J. and MeadoWatch volunteers collected the data; A.J. analyzed the data; A.J., E.J.T. and J.H.R.L. led the writing of the manuscript. All authors contributed critically to the drafts and gave final approval for publication.

## Data Availability Statement

All code will be made available on public GitHub repositories and will be archived in Zenodo.

## 6. Appendix A

### Mask R-CNN

We used Mask R-CNN (Mask Region based CNN), a deep learning algorithm that generates bounding boxes and segmentation for each instance of an object (here a flower) in an image (Fig. A.1). It is classified as a two-pass network; where the first part is dedicated to feature learning, while the second part is for detection and segmentation. In the end, the network identifies individual objects in an image and localizes each of them with a bounding box (Machefer et al. 2020) along with predicting pixel level segmentation masks. Instance segmentation in Mask R-CNN comprises of object detection and semantic segmentation. Mask R-CNN is an improvement upon Faster R-CNN (Ren et al. 2017) which used region proposal and classification networks to detect region of interest (RoI) and bounding boxes. Mask R-CNN supports various backbones as feature pyramid network (FPN) in order to produce feature maps of the input image. Region proposal network (RPN) is then applied atop feature maps that returns object proposals with corresponding objectness scores. Mask R-CNN identifies the anchor boxes based on the IoU, which is a critical step in the first part of the network (RPN). Objectness tells how good the model did to predict an object regardless of class. A RoIAlign layer is then applied to filter the proposals; the predecessor (Faster R-CNN (Ren et al., 2017)) used RoI pooling which made it slower. The key feature is the use of RoI Align instead of RoiPool (quantization free layer). The proposals are then passed to a fully connected layer that returns class and bounding boxes for the objects. Simultaneously, the mask branch that is a fully convolutional network (FCN) works on each RoI, and outputs a segmented mask at a pixel level. Mask R-CNN uses class loss, box loss and mask loss for scoring the model. In our study Mask R-CNN was configured with ResNext 101 backbone.

**Figure A1:**
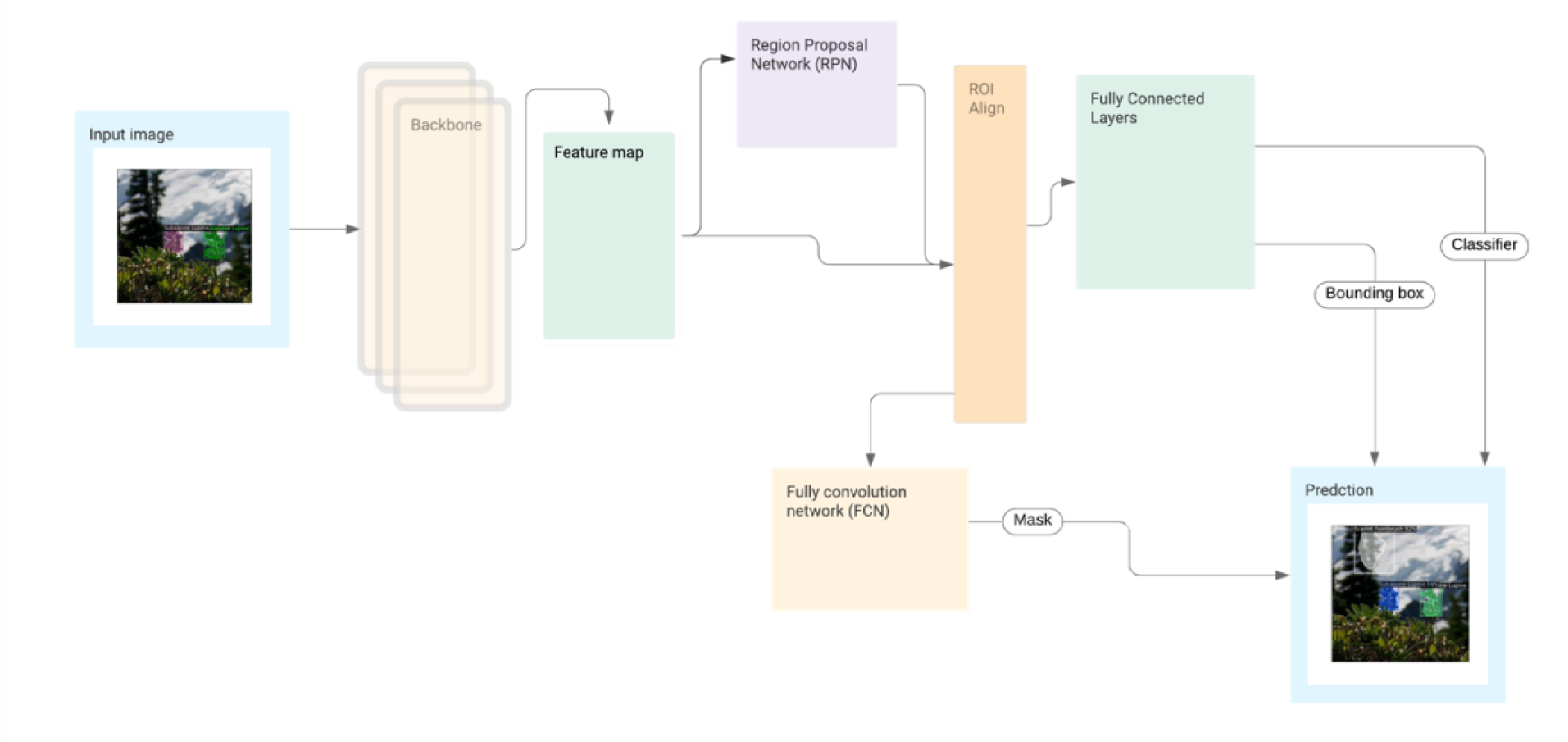
High level architecture of the Mask R-CNN algorithm.

### YOLO

YOLO stands for “You Only Look Once” that uses regression to achieve object detection and localization (Redmon et al. 2016). YOLO is a one stage method that works by regressing the bounding boxes and class probabilities and is known for its accuracy and high speed (Fig. A.2). YOLO has constantly evolved by releasing significant iterations with changes to its network structure resulting in better speed, consistent classification accuracy and recognizing smaller targets (Redmon et al. 2016, Redmon and Farhadi 2017, 2018). The YOLO method divides each input into regions or grids (S by S where S =8 for e.g.). For each grid it predicts B bounding boxes with confidence scores along with class conditional probabilities C. Confidence score translates to accuracy of the predicted object(s) in a grid. YOLO employs non-maximum suppression (NMS) similarly to Mask R-CNN if there are multiple candidate bounding boxes for the same target object. The loss function has three parts: objectness, box and classification. Objectness like in Mask R-CNN is a measure of how well the model did to predict an object irrespective of the class, Box tells how close the predicted box aligns with the actual box regardless of class and Classification loss tells how well each grid is classified with respect to its class probabilities. The YOLO algorithm penalizes the model’s inability to draw the bounding box correctly and where it missed estimating it completely. YOLO v5 was the last publicly available version by the original authors, and the recent release (unofficially known as Yolo v5) is now maintained by Ultralytics (GitHub handle - https://github.com/ultralytics/yolov5)

**Figure A2:**
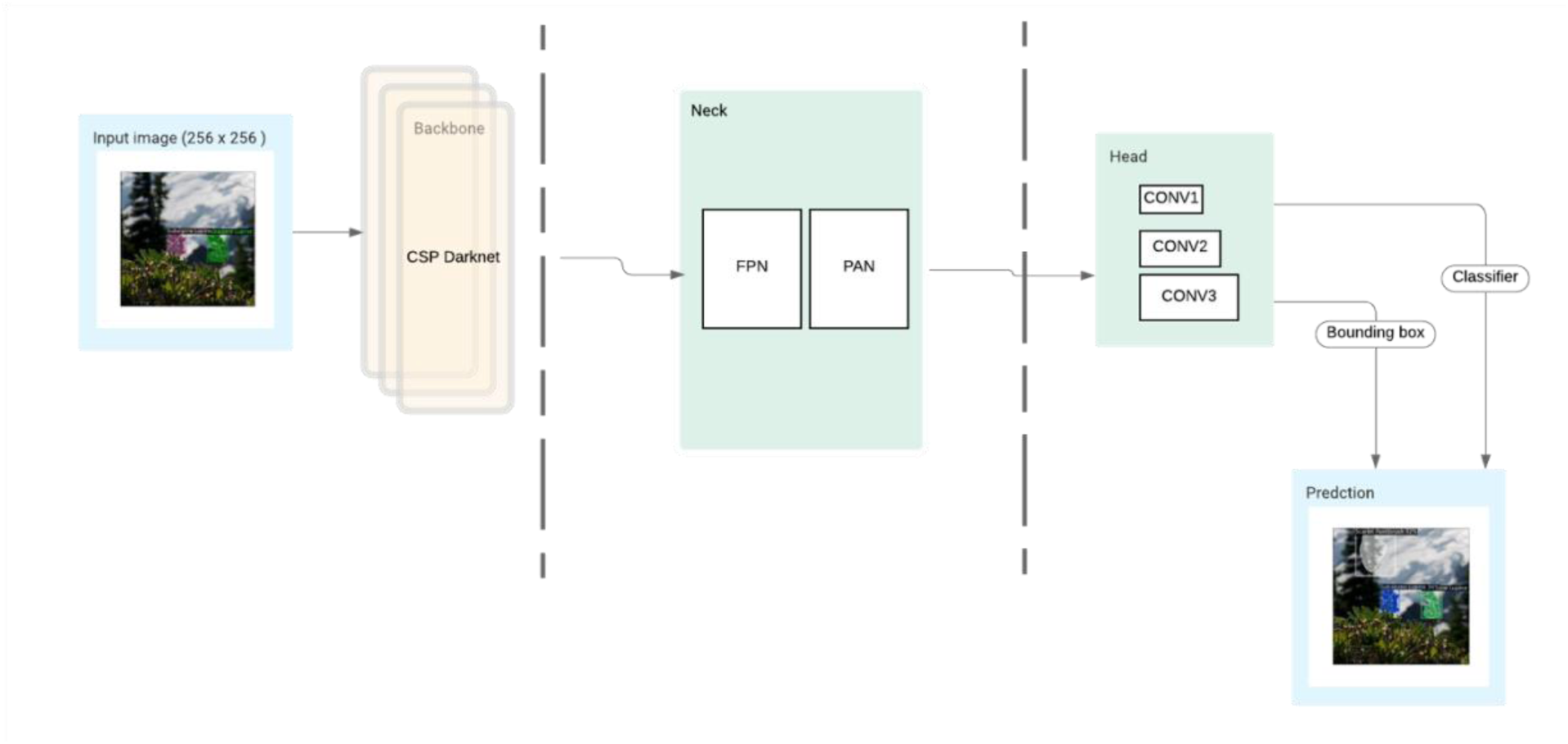
High level architecture of the YOLO algorithm.

### Retina-Net

RetinaNet is a three-part learner where the convolution layer learns from the architecture of the images (Lin et al. 2017, Li and Ren 2019). The architecture uses a feature pyramid network (FPN) backbone on the top of ResNet architecture to generate a rich, multi-scale convolutional feature pyramid (Fig. A.3). Two subnetworks are attached to it next – classification subnet and regression subnet. The first subnetwork is for classifying anchor boxes, while the second subnetwork is for regressing anchor boxes to the ground-truth boxes. The complex head and body of the network involves convolutions which reduces the image to a deep low-resolution image. The network is tuned by using the regression loss and the classification loss. Both metrics look at overall objectness and classification ability of the model. Regression loss is calculated by using smooth L1 whereas classification is evaluated by Focal loss.

**Figure A3:**
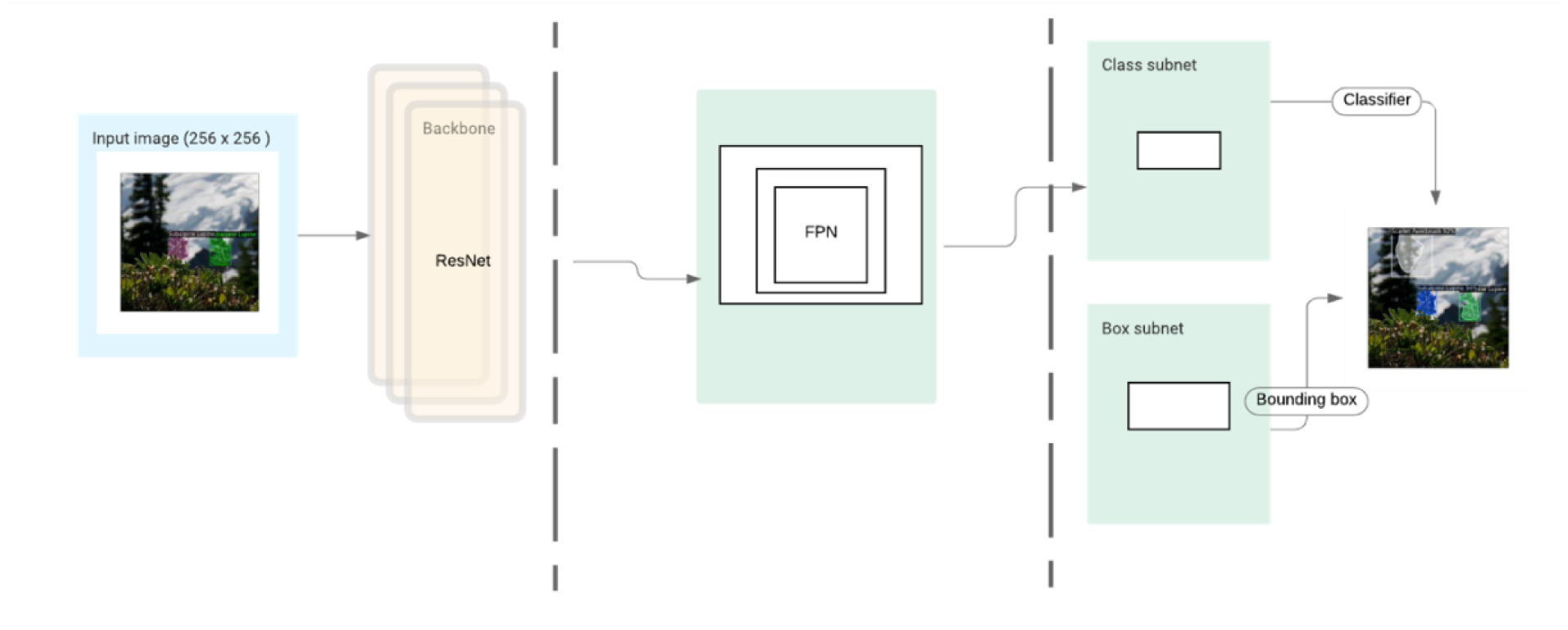
High level architecture of the RetinaNet algorithm.

## 7. Appendix B

### Predictions by RetinaNet and YOLO

**Figure B1:**
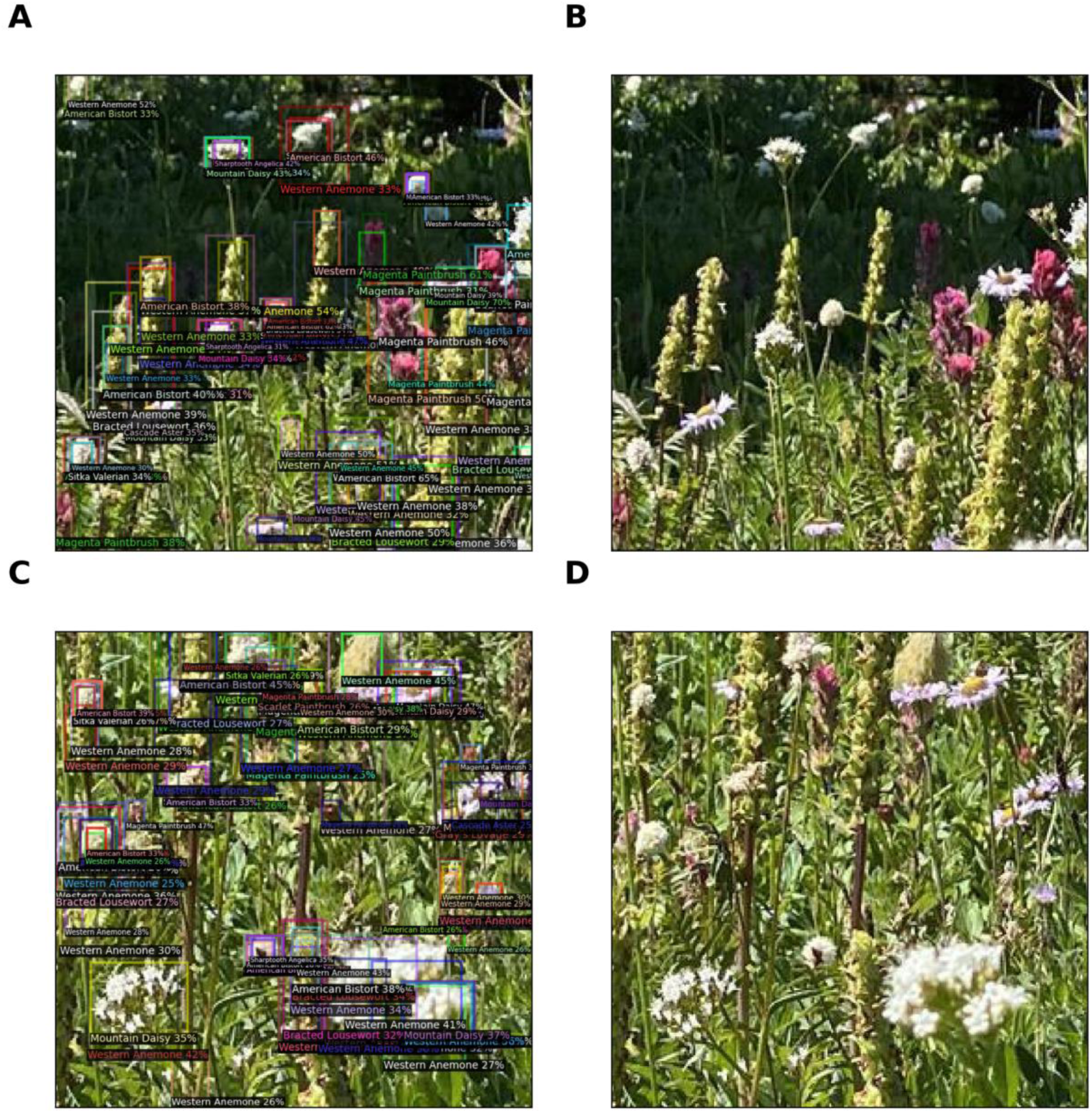
Predictions from the RetinaNet method showing the fluency of the model; the percent next to the predicted class represents model confidence. (A and B) Meadow in peak flowering with predictions (B) corresponding original tile. (C and D) Another meadow in peak flowering with predictions (D) corresponding to the original tile.

**Figure B2:**
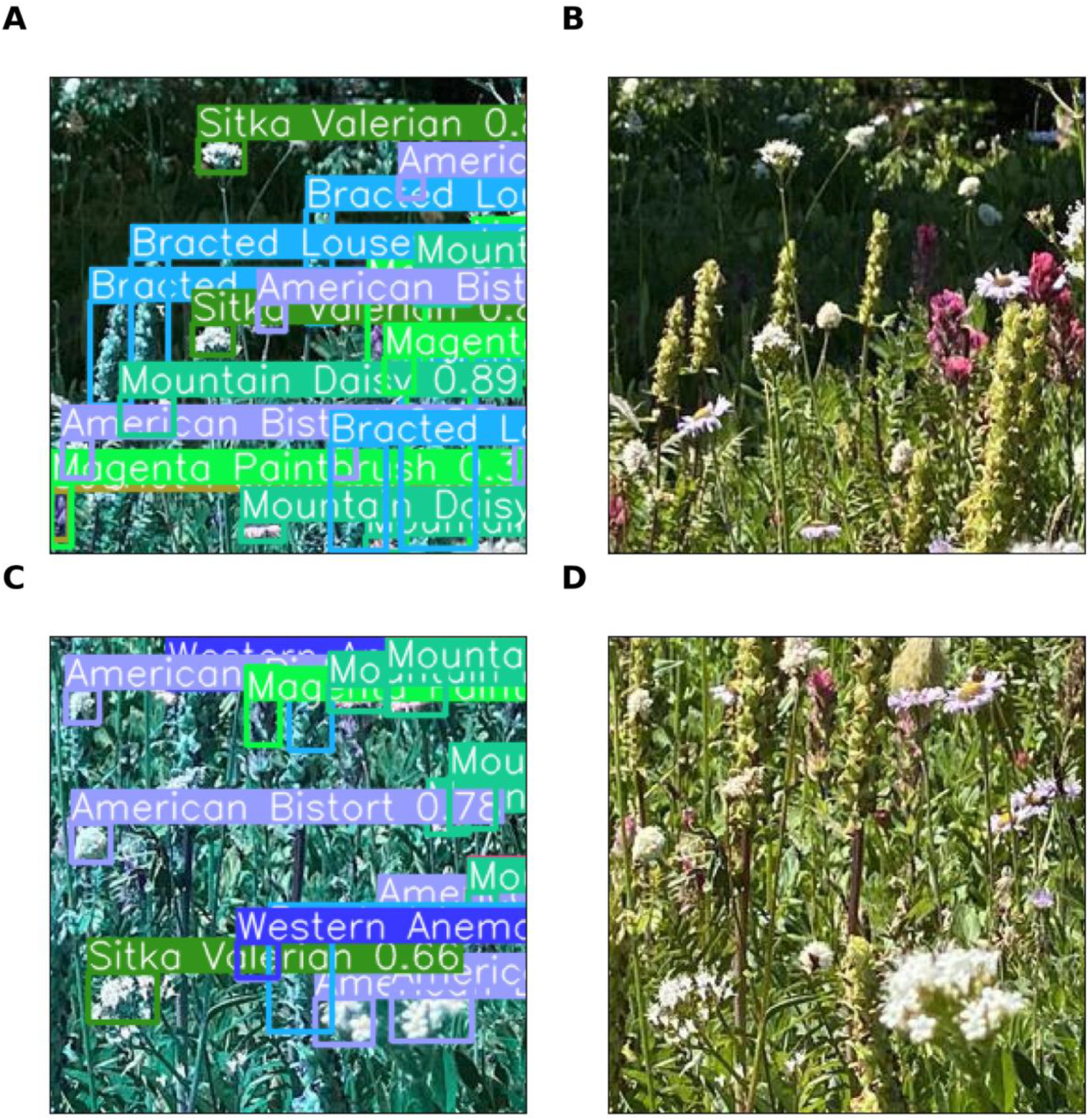
Predictions from the YOLO method showing the fluency of the model; the percent next to the predicted class represents model confidence. (A and B) Meadow in peak flowering with predictions (B) corresponding original tile. (C and D) Another meadow in peak flowering with predictions (D) corresponding to the original tile.

## 8. Appendix C

### Hyperparameters for the three methods

**Table C1:** Results of anchor box variation with RetinaNet. Used batch size of 10 and ran it for 250 epochs (iterations).

| Anchor Box Size | AP - Bounding Box |
| --- | --- |
| [4, 8, 16, 32, 64] | 6.99 |
| [8, 16, 32, 64, 128] | 9.05 |
| [16, 32, 64, 128, 256] | 13.41 |
| [32, 64, 128, 256, 512] | 8.22 |

## 9. Appendix D

### Correlation between richness as inferred by Mask R-CNN, YOLO and RetinaNet for August 2020 dataset

**Figure D1:**
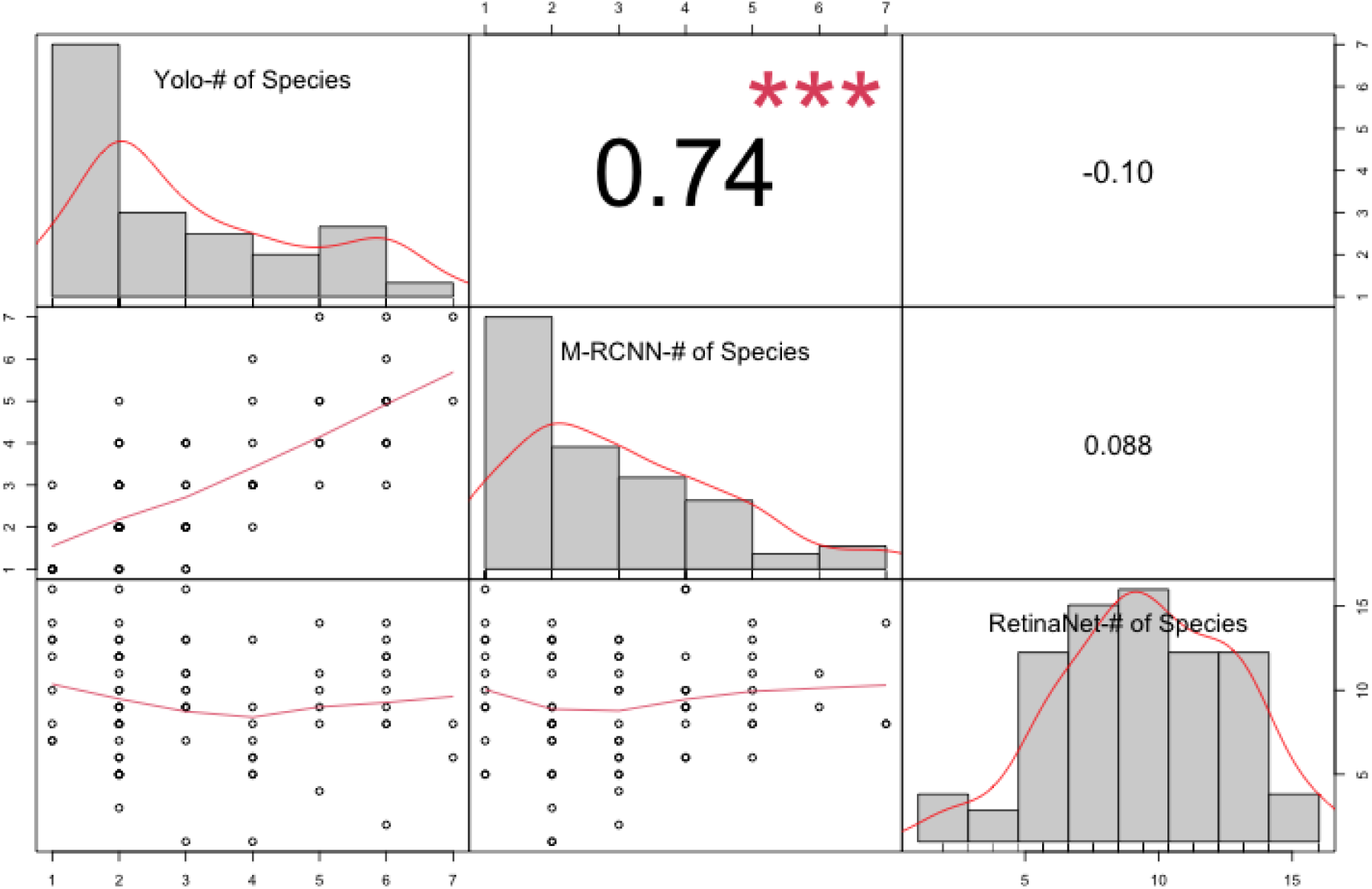
Correlation between richness (number of unique flowers) as calculated by Mask R-CNN, YOLO and RetinaNet for all the collections from August 2020. Significant association is found between YOLO and Mask R-CNN

## Notes

### Competing Interest Statement

The authors have declared no competing interest.

### Summary of Updates

Emphasized the significance of alpine meadows and our study's contribution to addressing existing gaps. Expanded upon the relevance of deep neural networks in the context of ecology and conservation.Introduced predictions from the other two methods (Yolo and RetinaNet) where applicable. Added domain specific metrics - species richness and Shannon-Wiener Index (a measure of diversity in species). Included comprehensive data on species abundance and richness derived from all three methods. Enhancements have been made to all figures to ensure improved clarity.

